# The role of feedback in responding to gradual and abrupt visuo-proprioceptive cue conflict

**DOI:** 10.1101/2024.09.12.612772

**Authors:** R. Babu, R. Matharu, C. Lo, H.J Block

## Abstract

When people observe conflicting visual and proprioceptive cues about their static hand position, visuo-proprioceptive recalibration results. Recalibration also occurs during gradual or abrupt visuomotor adaptation, in response to both the cue conflict and sensory prediction errors experienced as the hand reaches to a target. Here we asked whether creating a cue conflict gradually vs. abruptly, or providing error feedback, affects recalibration in a static hand. We examined participants’ responses to a 70 mm visuo-proprioceptive conflict, imposed by shifting the visual cue forward from the proprioceptive cue (static left hand). Participants pointed with their unseen right hand to indicate perceived bimodal and unimodal cue positions. Conflict was introduced gradually (groups 1 and 2) or abruptly (groups 3 and 4), with performance feedback present (groups 2 and 4) or absent (groups 1 and 3). For abrupt groups, most behavioral change occurred immediately after the conflict began. No-feedback groups (1 and 3) showed comparable magnitudes of overall recalibration, indicating that abrupt and gradual conflicts result in similar recalibration but with different timings. Motor adaptation was evident in the indicator hand with performance feedback (groups 2 and 4). However, performance on a static ruler task suggests proprioceptive recalibration also occurred despite the presence of feedback. Control groups confirmed accurate performance on the pointing task despite the visual cue shift. These findings highlight the distinct timing of recalibration mechanisms for gradual versus abrupt cue conflicts and potential smaller contribution of error mechanisms for a static conflict.

**New and Noteworthy:** The brain may handle spatial conflicts between visual and proprioceptive cues differently for a dynamic hand undergoing visuomotor adaptation than for a static hand. In a static hand, abrupt conflict triggered immediate recalibration without further adjustment, and feedback had little impact on recalibration. This suggests varying roles of multisensory and error mechanisms across motor contexts, underscoring the importance of examining a variety of motor contexts to understand and predict behavior.

## INTRODUCTION

When we try to use a computer mouse with an unfamiliar cursor gain, the brain adjusts hand movements to compensate for the unexpected cursor movements (sensory prediction errors). This process of visuomotor adaptation, commonly studied with a cursor rotation paradigm, is accompanied by shifts (recalibration) in the brain’s proprioceptive and visual estimates of hand position (Block & Bastian, 2011; van Beers et al., 2002). While visual recalibration is not often examined in this context, proprioceptive recalibration has some clear characteristics. The magnitude of proprioceptive recalibration is less than the motor adaption magnitude (Cressman & Henriques, 2009; Rand & Heuer, 2019; Redding & Wallace, 2006; Salomonczyk et al., 2011), and it occurs at a slower rate compared to motor adaptation (Zbib et al., 2016). This recalibration is robust and persists on removing visual feedback during (Barkley et al., 2014), or at the end of movement (Maksimovic et al., 2020), and on increasing explicit awareness of the perturbation (Modchalingam et al., 2019).

Importantly, perceptual recalibration occurs even in the absence of movement, simply in response to a spatial conflict between visual and proprioceptive cues of hand position. With sufficient exposure, both visual and proprioceptive estimates recalibrate in response to the mismatch (Block & Bastian, 2011; Hay & Pick Jr., 1966; Rand & Heuer, 2019), though there are other studies that only find proprioceptive recalibration (Simani et al., 2007; Zbib et al., 2016).

Recalibration in response to a cue conflict differs in some ways from the recalibration associated with visuomotor adaptation, which is thought to arise from both sensory prediction errors and cue conflict (Rossi et al., 2021). Proprioceptive recalibration observed in visuomotor adaptation studies develops quickly and persists (Ruttle et al., 2021, 2022) Visuo-proprioceptive recalibration is also robustly retained in cue-conflict studies (Wali et al., 2023). In a visuomotor adaptation study, the cursor perturbation may be applied abruptly or gradually; the magnitude of proprioceptive recalibration does not appear to differ, provided the same final distortion (e.g., a 30° cursor rotation) was used. (Cressman & Henriques, 2009; Salomonczyk et al., 2011). To our knowledge, recalibration in response to abrupt vs. gradual visuo-proprioceptive cue conflict alone has not previously been tested. However, visuo-proprioceptive recalibration in a cue-conflict study has been observed to be higher in magnitude for gradual mismatches that occur more rapidly than for gradual mismatches that occur slowly (Babu et al., 2023). A critical aspect of this recalibration is that when the cue conflict is maintained at a constant level across trials, no further recalibration occurs. In contrast, recalibration in a motor adaptation paradigm continues to increase even if the perturbation is maintained at a constant level, until till around 50% of the perturbation is compensated for (Cressman & Henriques, 2009, 2010).

In natural behavior, all these processes likely coexist and interact to some degree, making it challenging to know which mechanism makes what contribution to behavior and learning. In the research setting it is possible to isolate the mechanisms to some degree, for example by removing motor error information so that only cue conflict drives recalibration, or by using a force perturbation instead of a cursor rotation so that motor adaptation and recalibration in response to sensory prediction errors can be examined in the absence of a cue conflict. A limitation of this body of work is that it has been done largely using a single paradigm: cursor rotation, where a cursor’s movement direction is gradually or abruptly rotated from the unseen hand’s movement direction by an angular offset while participants make center-out reaches to visual targets (Cressman & Henriques, 2009; Redding & Wallace, 2006; Salomonczyk et al., 2011). Variations of this task have yielded major advances in our understanding of sensorimotor control, but in the case of proprioceptive and visual recalibration, such findings must be reconciled with the cue conflict literature, which uses other tasks.

When motor adaptation is not the goal, perceived hand position is commonly measured and/or perturbed with a bimanual task, using an “indicator” hand to indicate the person’s perception of the “target” hand’s position when visual, proprioceptive, or both types of information about the stationary target are available (Smeets et al., 2006; van Beers et al., 1996, 1998, 1999, 2002). For bimodal targets, the visual and proprioceptive cues can be presented veridically or mismatched. Unlike a cursor rotation, this task does not impose a cue conflict in the context of making a reaching movement to a visual target, where the conflict increases the further one moves from the home position. Instead, the proprioceptive and visual cues are themselves the targets, with the conflict presented in bimodal cues as a static offset.

Here we used the bimanual cue conflict task to examine the effect on recalibration of two factors commonly associated with visuomotor adaptation: perturbation type (abrupt or gradual) and error feedback (present or absent). Participants pointed with their unseen right hand to indicate perceived cue positions in a repeating sequence of bimodal and unimodal targets. A 70 mm conflict was introduced gradually (groups 1 and 2) or abruptly (groups 3 and 4) by shifting the visual cue forward. Based on the finding that among gradual cue conflicts, recalibration is greater for more rapidly-imposed conflicts (Babu et al., 2023), we might predict even higher recalibration for an abrupt mismatch. We might also predict that any recalibration occurs immediately for an abrupt mismatch, with no additional recalibration while the cue conflict is held constant (Babu et al., 2023).

Indicator finger performance feedback was present (groups 2 and 4) or absent (groups 1 and 3). Because we expected performance feedback to cause motor adaptation in the indicator hand for groups 2 and 4, indicator finger endpoints do not solely reflect visual and proprioceptive recalibration in the target estimates in these groups; we therefore also assessed proprioceptive recalibration with a ruler estimation task that did not involve the indicator hand at all. We predicted that indicator finger feedback would provide sensory prediction errors that, combined with the cue conflict, might drive higher proprioceptive recalibration as seen in the ruler task. Finally, groups 5 and 6 performed the task with unimodal cues only (group 5) or visual cues only (group 6) to control for any biasing effect of displacing the visual cue away from the subject and potentially making the indicator finger movements more onerous.

## METHODS

### Participants

A total of 96 individuals participated in the study (42 male, mean age 24.09). Participants were randomly assigned to one of six groups (N=18 for groups 1-4 and N= 16 for groups 5 and 6). All participants were right-handed (85.04 mean score on the Edinburgh handedness questionnaire (Oldfield, 1971), young adults (18-45 years old) with normal or corrected to normal vision. They reported no known neurological diseases, attention disorders, or musculoskeletal injuries. The protocols were approved by the Institutional Review Board (IRB) of Indiana University Bloomington and were performed in accordance with the IRB guidelines and regulations.

### Session and groups

Participants completed one session lasting approximately 1.5 hours. Sessions began by introducing the participant to the experiment through instructional PowerPoint slides. For groups 1-4, this was followed by two tasks performed at the same apparatus: the pointing task and the ruler task, described below. For these groups, the pointing task differed during Block 2, when a 70 mm visuo-proprioceptive mismatch was imposed gradually (groups 1 and 2) or abruptly (groups 3 and 4) and performance feedback was present (groups 2 and 4) or absent (groups 1 and 3) (Fig. 1A-B). Groups 5 and 6 had only a pointing task.

**Fig 1.**
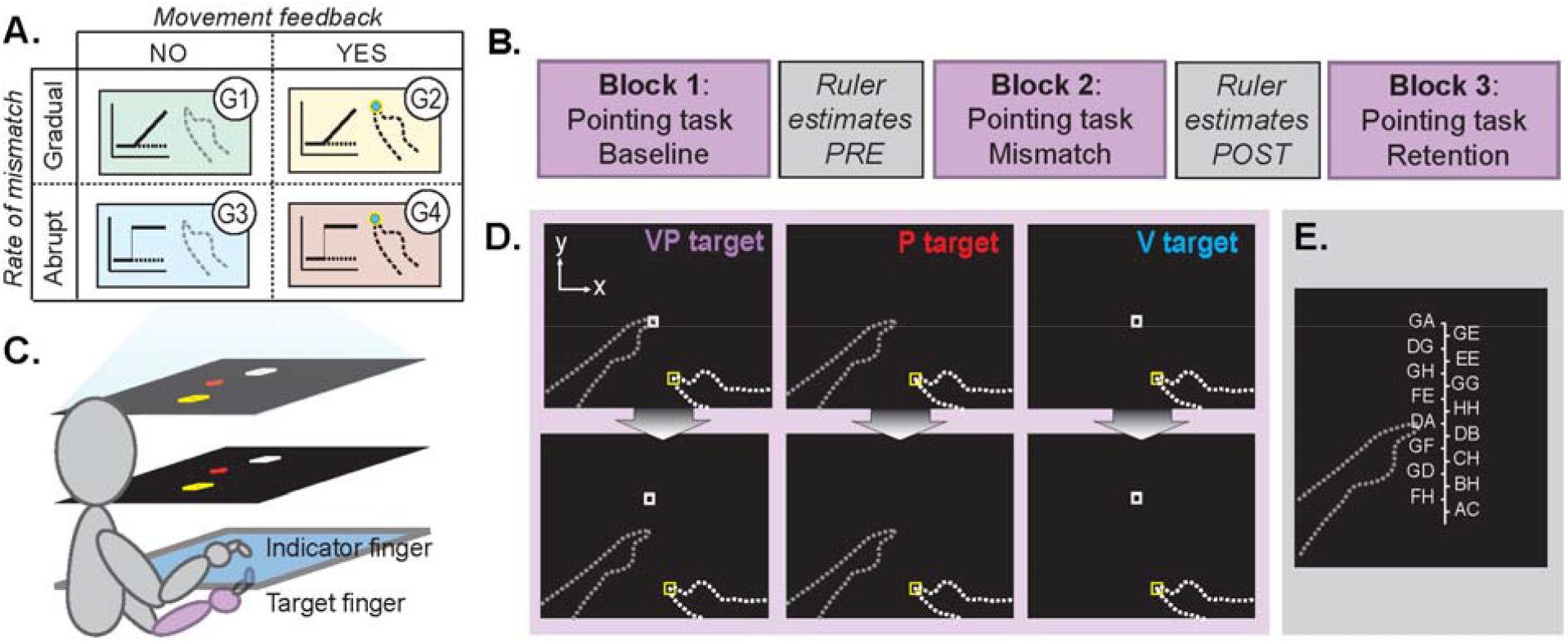
Experimental groups and methods. **A**. Groups 1-4 design: Participants were randomly assigned to their respective groups, which differed in Block 2 of the pointing task. The visuo-proprioceptive mismatch in Block 2 occurred gradually for groups 1 and 2 and abruptly for groups 3 and 4. Groups 1 and 3 received no feedback about their performance at any time, while groups 2 and 4 received feedback about their performance at estimating bimodal targets during Block 2 of the pointing task. **B**. Session design: Participants performed three blocks of the pointing task, with the ruler task occurring before and after Block 2. The visuo-proprioceptive mismatch was introduced in Block 2. **C**. Task apparatus: The task display was projected onto a screen (top layer) and viewed on a mirror (middle). Participants perceived the display to be in the same plane as the touchscreens (bottom layer). **D**. Pointing task targets: VP targets were used to induce a visuo-proprioceptive mismatch, where the white box was initially on top of the target finger (top box), but later shifted forward a maximum of 70mm from the target finger (bottom box). Dashed lines not visible to subjects. **E**. Ruler estimates task: Participants viewed a “ruler” of letter codes and reported the code they perceived to be over their unseen target finger. The letter codes changed between each trial.

### Apparatus

Participants were seated at a 2D virtual reality apparatus that used a double-sided infrared touchscreen (PQ Labs) with an area of 75cm x 100 cm, a mirror positioned above the touchscreens and below eye-level, and a projection screen above eye level (Fig. 1C). With this set-up, the visual display appeared to subjects to be in the same horizontal plane as the touchscreens. No direct vision of the hands was possible during the task. The right “indicator” hand remained on top of the touchscreen layer at all times, while the left “target” hand remained below the touchscreen layer. Their upper arms and shoulders were occluded using a black drape.

### Pointing task

#### Targets

Without vision of the hands, participants were instructed to use their right indicator finger to point on the touchscreen to the perceived location of three different types of targets: visual only (V), proprioceptive only (P), and combined (VP) (Fig 1D). The visual target was a 1cm white square projected in the task display. The proprioceptive target was the participant’s target index fingertip that was positioned on either a rough or smooth tactile marker on the bottom of the touchscreen. The combined VP target was indicated by a white box projected on top of the target finger’s location. Participants were explicitly informed that the white box was positioned on top of the left index finger for VP trials, although this was the case only at the beginning of the task.

#### Single-trial procedure

At the start of each trial, participants were instructed to place their unseen right “indicator” index finger in the center of a 1cm yellow box that appeared in any of five positions on the touchscreen apparatus. Their finger placement was guided by a 0.8 cm blue dot which disappeared once the participant’s finger aligned with the yellow box. Once in the starting position, participants were cued via audio instructions to keep their eyes on a red cross that appeared at random coordinates within 10 cm of the target position. Participants were then told to place their left “target” index finger on the smooth or rough tactile marker beneath the touchscreen surface for P and VP trials. For V trials, subjects were instructed to keep their target hand lowered.

Once both hands were correctly positioned, they heard a go signal (beep) and were instructed to place their indicator finger where they perceived the target to be. During practice reaches at the beginning of the session, subjects were asked to prioritize accuracy over speed and to avoid dragging their indicator finger along the glass. Once they felt the correct position was achieved, they held still for 2s after which the final target estimate was recorded and the trial ended with no feedback about performance accuracy, with limited exceptions: For groups 2 and 4 subjects who received performance feedback during Block 2 after VP trials only, the 0.8 cm blue dot appeared briefly in the location of the participant’s target estimate before disappearing prior to the start of the next trial (Fig. 1A).

#### Pointing task blocks

For groups 1-4, Block 1 (baseline) consisted of 15 V, 15 P, and 15 VP trials in pseudorandom order for groups 1-5 (Fig. 1B). VP trials were veridical. Block 2 (mismatch) consisted of 21 V, 21 P, and 42 VP trials in the repeating order V-VP-P-VP. During this block, the visual cue shifted forward relative to the proprioceptive cue either gradually, 1.67 mm at a time (groups 1 and 2) or abruptly (groups 3 and 4). Block 3 (retention) consisted of an additional 15 V and 15 P trials in pseudorandom order. The visual cue continued at its 70 mm forward location to avoid a sudden change that subjects might notice.

The purpose of group 5 was to assess whether unimodal target estimates would change in the absence of bimodal trials containing the cue conflict. Subjects in Group 5 experienced the same target sequence as Group 1, except that VP trials were omitted from Block 2 entirely. Thus, while the visual cue shifted forward at the same pace in both groups, it was never presented simultaneously with the proprioceptive cue for Group 5.

The purpose of group 6 was to assess whether unimodal V target estimates would shift relative to the actual target position in the absence of P or VP targets to drive visual recalibration. Group 6 participants therefore experienced only V targets in all 3 blocks. During Block 2, the V target was gradually shifted forward to achieve a total 70 mm shift after 84 trials, the same rate as groups 1 and 5.

After Block 3, group 1-5 participants were asked if they felt like the white square was always on top of their target finger, and if not, which direction and how much did they feel like it was off. Participants were also asked to rate their quality of sleep, level of attention, and level of fatigue they felt from the experiment on a scale of 1 to 10.

#### Target endpoint shift

For proprioceptive and visual estimates separately, four comparisons were made between different pairs of timepoints in the task: Initial response to the perturbation (late Block 1 to early Block 2), change in performance during perturbation (early Block 2 to late Block 2), overall response to perturbation (late Block 1 to late Block 2), and forgetting (late Block 2 to late Block 3). For each of these comparisons, the y-dimension shift in target position estimate (e.g., proprioceptive shift ΔY_P_) was obtained by subtracting the mean y-coordinate on the first or last four trials of a given modality in the given block(s):

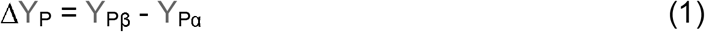

For example, for the proprioceptive initial response to the perturbation, Y_Pβ_ would be the mean of the first four P target estimates in Block 2, and Y_Pα_ would be the mean of the last four P target estimates in Block 1.

Visual endpoint shift (ΔY_V_) was computed similarly, except subtracted from 70 when comparing across timepoints that involved a 70 mm V target shift so the shift would be relative to V target position. Both visual and proprioceptive estimate shifts in the expected direction (closing the imposed gap) were thus set as positive values. The sum of proprioceptive and visual endpoint shift accounts for the total endpoint shift.

### Ruler estimation task

To quantify proprioceptive recalibration without using the indicator finger to point, subjects in groups 1-4 performed a proprioceptive estimation task using a ruler. The task consisted of 10 trials immediately before and 10 trials immediately after Block 2 of the pointing task (Fig. 1B). All 20 trials were performed with the left target index finger positioned on the rough tactile marker. The task display consisted of a 16-cm-long “ruler” displayed over the unseen target finger. To avoid numerical biases, each 0.5 cm along the ruler was labelled with a random two-letter code rather than a number. In each trial, participants verbally reported which two-letter code felt the closest to being directly on top of their left finger (Fig 1E). For each participant, the mean estimated position on the ruler was subtracted post minus pre to assess proprioceptive recalibration.

### Statistical analysis

Visual, proprioceptive and total recalibration at different stages was compared for the four groups using a 1-way ANOVA. Tukey post-hoc tests were performed to examine significant ANOVA effects. All hypothesis tests were performed two-sided with α = 0.05. Shapiro–Wilk’s tests revealed that the group data were normally distributed. Post-hoc tests with Bonferroni adjustments were performed on significant interactions.

A mixed method ANOVA with Task (Ruler, Pointing) x Session (Pre, Post) x Group was performed on the proprioceptive recalibration to understand how the different methods compare while measuring recalibration. Paired sample t-tests were used to compare the proprioceptive recalibration measured using the ruler task and computed from the reaching task for each group.

In experiment 2, along with the one-way ANOVA on visual recalibration we also performed a one-sample *t* test on each group’s raw recalibration against a test value of 0.

The data and analysis code used in this manuscript are available at https://osf.io/486ut/

## RESULTS

Subjects showed a variety of responses to the visuo-proprioceptive mismatch pointing task (Fig. 2). Block 1 performance appears similar across groups, as no mismatch or feedback was introduced yet. Different patterns are evident in Block 2. Generally, if feedback was provided about indicator finger endpoints on bimodal VP targets (G2 and G4), the amount of visual target undershoot decreased and proprioceptive target overshoot increased (Fig. 2). This may reflect a systematic change in motor commands to the indicator hand (motor adaptation) to compensate for undershooting the visual component of the VP target as the mismatch occurs. Another pattern that emerges is that for the abrupt groups (G3 and G4), most of the change in target over/undershoot appears to have happened quickly, as soon as the 70 mm mismatch was introduced in Block 2.

**Fig 2.**
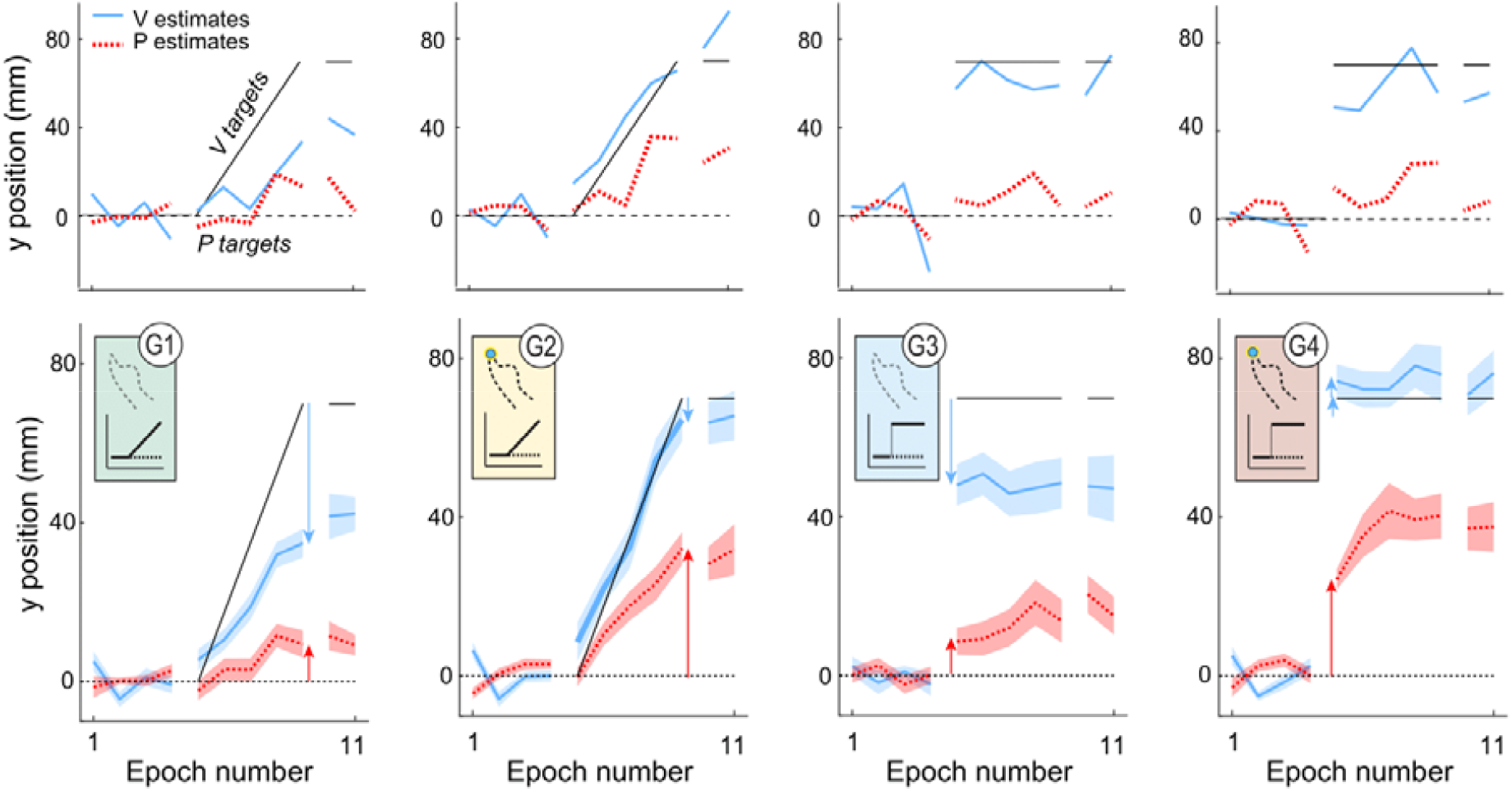
**Top row:** Example participants’ y-dimension indicator finger endpoints on P (red) and V (blue) targets, averaged every 4 trials for all 4 groups. The example subjects show varying amounts of change in visual (Δ*Y*_*V*_) and proprioceptive (Δ*Y*_*P*_) target estimates. **Bottom row:** Mean and SEM of performance for G1-G4, averaged every 4 trials. Irrespective of the mismatch condition, groups with feedback (G2 and G4) showed larger proprioceptive target overshoot but smaller visual recalibration undershoot compared to the corresponding groups with no feedback. Subjects in the abrupt mismatch groups (G3 and G4) responded immediately to the mismatch and mostly maintained it throughout the block while those in the gradual groups (G1 and G2) gradually changed their behavior as the mismatch progressed.

We compared 4 pairs of timepoints to analyze change in performance on visual and proprioceptive targets: the initial response to perturbation (early Block 2 minus late Block 1), change in performance during perturbation (late Block 2 minus early Block 2), overall response to perturbation (late Block 2 minus late Block 1), forgetting (late Block 3 minus late Block 2).

### Initial response to perturbation

The initial response to perturbation was calculated as the difference in mean visual or proprioceptive end points at the beginning of block 2 (when the perturbation was first imposed) and that at the end of block 1. A one-way ANOVA suggests a significant difference in the initial response to the perturbation among groups (F (3,68) = 4.119, p = 0.010) for visual target performance. The Tukey’s Post Hoc comparison showed that Group 3 was significantly different from Group 4 (p = 0.005) (Fig 3A).

**Fig 3.**
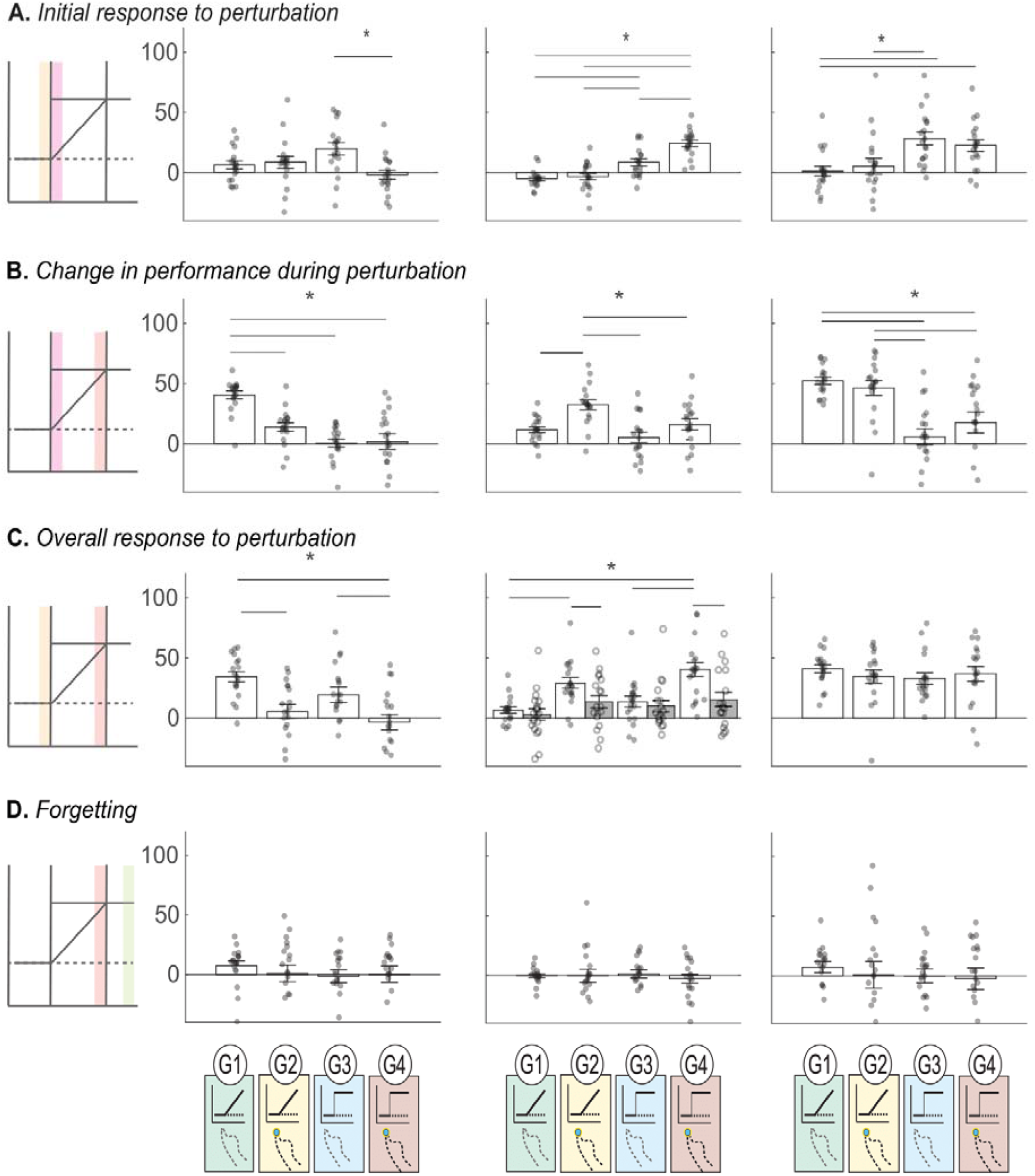
Group changes in performance on visual targets (left column), proprioceptive targets (middle column), and sum of both (right column). Means and SEM with dots representing individual participants. *p<0.05. **(A)** Initial response to perturbation for the 4 groups which is calculated as the difference between the beginning of Block 2 and end of Block 1. **(B)** Change in performance during perturbation which is response over Block 2 and is calculated as the difference between end of Block 2 and beginning of Block 1. **(C)** Overall response to the perturbation which is calculated as the difference from the end of Block 1 to the end of Block 2. For proprioception, the ruler task (grey bars) and pointing task (white bars) are compared side-by-side. **(D)** Forgetting which is the difference between late Block 2 to late Block 3.

There was also a significant effect of group on proprioceptive target initial response (F (3,68) = 27.244, p < 0.001). The post-hoc analysis showed that groups 1 and 3 (p = 0.002), groups 1 and 4 (p < 0.001), groups 2 and 3 (p = 0.011), groups 2 and 4 (p < 0.001), and groups 3 and 4 (p < 0.001), all varied with respect to the initial response to perturbation (Fig 3A).

The total endpoint shift (sum of visual and proprioceptive target changes) as a result of the initial perturbation showed a significant effect of group response (F (3,68) = 5.996, p = 0.001). The post-hoc analysis showed that groups 1 and 3 (p = 0.004), groups 1 and 4 (p = 0.032) and groups 2 and 3 (p = 0.017) varied with respect to the initial response to perturbation (Fig 3A).

### Change in performance during perturbation

Change in performance during the perturbation was determined as the difference in mean endpoint y-coordinates between the beginning and end of Block 2. There was a significant effect of group on visual target performance change (F (3,68) = 18.844, p < 0.001). Post-hoc comparisons revealed that Group 1 differed significantly from 2 (p < 0.001), 3 (p < 0.001), and 4 (p < 0.001), suggesting end point feedback and perturbation rate both affected visual target response (Fig 3B).

There was also a significant group effect on proprioceptive target response over the course of Block 2 (F (3,68) = 8.630, p < 0.001). Specifically, Groups 1 and 2 (p = 0.002), Groups 2 and 3 (p < 0.001), and Groups 2 and 4 (p = 0.024) all differed with respect to proprioceptive targets during perturbation (Fig 3B).

The total endpoint shift as a result of the perturbation was also significantly different across groups (F (3,68) = 12.072, p < 0.001). Specifically, Group 1 differed from Groups 3 (p < 0.001), and 4 (p = 0.001), and Group 2 differed from Groups 3 (p < 0.001) and 4 (p = 0.016) as per the post-hoc analysis (Fig 3B).

### Overall response to perturbation

To account for the differences in groups with abrupt and gradual perturbations, overall visual target performance change in response to the perturbation was defined as the difference in indicator finger endpoints on visual targets at the end of Block 2 (end of perturbation) compared to the end of Block 1 (before perturbation). The data suggests a significant difference between overall visual target change across groups (F (3,68) = 8.889, p < 0.001). In the post-hoc comparison, Groups 1 and 2 (p = 0.002), Groups 1 and 4 (p < 0.001), and Groups 3 and 4 (p = 0.035), were significantly different (Fig 3C).

Overall change in performance in proprioception included both the pointing task with proprioceptive targets from late Block 1 to late Block 2 timepoints, and the ruler task that was performed pre- and post-Block 2. The mixed effects model of Task (Ruler vs Pointing) X Group X Timepoint showed main effects of task (F (1,68) = 412.983, p<0.001) and timepoint (F (1,68) = 70.825, p<0.001), but no main effect of group (F (3,68) = 2.265, p = 0.089). There were also Timepoint X Group (F (3,68) = 6.828, p<0.001), and Timepoint X Task (F (1,68) = 17.363, p<0.001) and Task X Group X Timepoint (F (3,68) = 3.223, p = 0.028) interaction effects. There was no Task X Group (F (3,68) = 2.313, p = 0.084) interaction effects. This suggests that performance in proprioceptive estimates changed differently across groups between the two timepoints, but this difference was related to whether ruler or pointing task is considered.

For the pointing task, there was a significant group difference in proprioceptive target pointing at the end of Block 2 with respect to the end of Block 1 (F (3,68) = 11.421, p < 0.001). Specifically, Groups 1 and 2 (p = 0.004), Groups 1 and 4 (p < 0.001), and Groups 3 and 4 (p < 0.001) differed from each other (Fig 3C).

Paired sample t-tests were used to compare within each group the effect of Task (pointing vs. ruler). The effect of Task was significant only for the groups with feedback – Groups 2 (t (17) = 2.381, p= 0.029) and 4 (t (17) = 4.70, p< 0.001)—while it was not significant for the groups without feedback, Groups 1 (t (17) = 0.788, p= 0.442) and 3 (t (17) = 0.59, p= 0.563).

For the ruler task, there was a main effect of time (F (1,68) = 16.328, p< 0.001) but no main effect of group (F (3,68) = 0.801, p= 0.498) and no significant difference across groups between the post and pre-measure (F (3,68) = 1.134, p= 0.341). Also, on performing a paired t-test comparing pre vs. post in each group, Group 2 (t (17) = -2.6614, p= 0.016) and Group 4 (t(17) = -2.614, p= 0.018) showed significant difference between the pre and post values while Group 1 (t(17) = -0.605, p= 0.553) and 3 (t(17) = -2.061, p= 0.055) did not show any significant difference.

There was no significant group difference in total endpoint shift at the end of Block 2 with respect to the end of Block 1 (F (3,68) = 0.721, p = 0.543) (Fig 3C).

### Forgetting

Forgetting was calculated as the difference between late Block 3 and late Block 1 for both vision and proprioception. The one-way ANOVA across all groups showed no significant differences between the groups for visual targets (F (3,68) = 0.592, p= 0.623) and no significant difference for proprioceptive targets (F (3,68) = 0.192, p = 0.901)(Fig 3D). There were also no group differences in the total forgetting (F (3,68) = 0.366, p= 0.778).

Group 1-4 participants were questioned at the end of the session about whether they perceived the visual and proprioceptive components of the VP target as veridically aligned throughout the experiment. 5 subjects in Group 1 reported perceiving any degree of forward offset (the real offset direction), but twice as many subjects in Groups 2-4 reported perceiving a forward offset (Table 1). Subjects in all groups rated their level of attention and fatigue similarly, suggesting these factors are unlikely to account for group differences (Table 1).

**Table 1.** Summary of participant ratings and perception responses (means and SEM).

|  | **Group 1** | **Group 2** | **Group 3** | **Group 4** | **Group 5** | **Group 6** |
| --- | --- | --- | --- | --- | --- | --- |
| Fatigue | 3.67 ± 0.40 | 4.17 ± 0.51 | 3.89 ± 0.35 | 4.11 ± 0.50 | 4.94 ± 0.47 | 4.06 ± 0.45 |
| Attention | 7.81 ± 0.36 | 7.39 ± 0.34 | 7.33 ± 0.34 | 7.533 ± 0.24 | 7.31 ± 0.22 | 7.25 ± 0.27 |
| Reported forward offset magnitude (cm) | 0.50 ± 0.31 | 1.02 ± 0.34 | 1.98 ± 0.43 | 0.44 ± 0.25 | n/a | n/a |
| Subjects who reported forward offset | 3 | 8 | 8 | 3 | n/a | n/a |

### Visual recalibration and effort minimization: Control groups

To control for the possibility that undershooting the V target reflects some minimization of effort (resistance to moving the indicator finger far from the body), we compared Group 1 performance with two control groups: Group 5 had V and P trials but no VP trials in Block 2, while Group 6 had only V trials throughout the three blocks. All three groups had a gradual 70 mm forward shift of the V target during Block 2. Subjects in Group 1 showed both visual (42.27 mm) and proprioceptive (11.66 mm) endpoint shifts in Block 2, each of which was significantly different from zero (t(17) = 11.04, p<0.001 and t(17) = 4.61, p<0.001 respectively). Group 5 subjects showed nonsignificant proprioceptive (7.85 mm, t(15) = 1.797, p = 0.092) and significant visual endpoint shifts (15.74 mm, t(15) = 2.781, p = 0.014, Fig 5B). Group 6 subjects showed no significant visual endpoint shift (1.981 mm, t(15) = 0.295, p = 0.772). The visual endpoint shift differed significantly across the three groups (F (2,47) = 14.683, p <0.001). Post-hoc tests showed that Groups 1 and 5 (p = 0.003) and Groups 1 and 6 (p <0.001) differed significantly from each other (Fig 4C).

**Fig 4.**
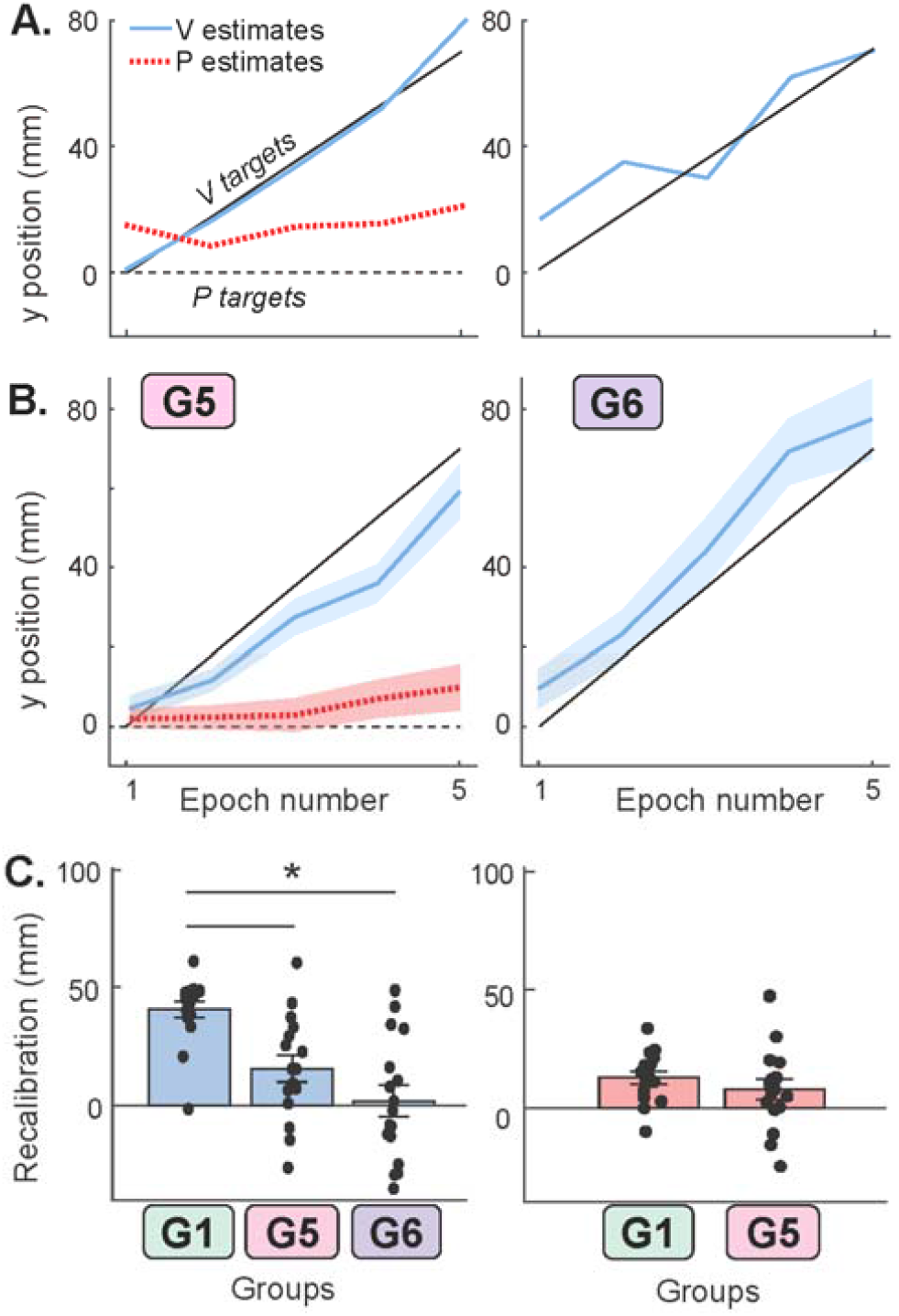
Control results. **(A)** End-point indicator finger position for example subjects in control groups 5 (left) and 6 (right). Like group 1, groups 5 and 6 experienced a gradual 70 mm forward shift of the visual target during block 2, with no indicator finger feedback. Group 5 had only V and P trials, and Group 6 had only V trials. **(B)** Group data showing mean and SEM indicator finger end-point positions. **(C)** Left: Mean and SEM of shift in V target endpoints (mm) of groups 1, 5, and 6. Group 1, which had V, P, and VP trials, showed greater increase in undershooting of V targets than Group 5, which had only V and P trials, and Group 6, which had only V trials. Only Group 1 and 5 differed significantly from zero. Right: Mean and SEM of shift in P target endpoints (mm) of groups 1 and 5. This parameter was significantly different from zero for Group 1, but not different from Group 5.

Participants were asked to rate their fatigue and attention on a scale of 1-10 at the end of the session, with 10 being maximum fatigue or attention. Participants were also asked about VP target perception: “Did it feel like the white square was always on top of your finger, or did it sometimes feel off?” Participants who reported that it felt off were asked follow-up questions to find out the perceived offset direction and maximum amplitude.

## DISCUSSION

The current study aimed to clarify the nature of recalibration in response to conflicting cues about static hand position. We first examined the effect of abrupt vs gradual perturbation and the effect of endpoint error feedback using a 2×2 design. While the abrupt groups showed an immediate effect of the cue-conflict, the overall response did not differ from the gradual groups. Motor adaptation was evident in the indicator hand with performance feedback, but the static ruler task suggests proprioceptive recalibration still occurred in the target hand. Finally, findings from the control groups indicate that subjects point accurately even when the visual target gradually moves 70mm away.

### Effect of abrupt and gradual mismatch on visuo-proprioceptive recalibration: Group 1 vs. 3

We previously found that magnitude of recalibration is greater when the rate of a gradually-imposed mismatch is faster; however, complete compensation (i.e., 70 mm total recalibration in response to a 70 mm cue conflict) was not achieved with any of the mismatch rates tested (Babu et. al, 2023). We hypothesized that the abrupt mismatch in the present study would result in greater recalibration than the gradual condition, as an abrupt mismatch may be more similar to how cue conflicts are experienced in real life.

For example, wearing bifocals is a quick abrupt change with the two different magnifications but something that we can quickly adjust to. However, total endpoint shift in the gradual and abrupt groups with no feedback (1 and 3) was similar in magnitude; the gradual group did most of this shift over the course of the perturbation, while the abrupt group had the full shift happen when the perturbation first started. Neither group compensated completely for the 70 mm offset, but instead recalibrated consistently with previous studies (Babu et al., 2023; Hsiao et al., 2022). Likewise, in the visuomotor rotation paradigm the resultant proprioceptive recalibration has been reported to be ∼50% of the motor adaptation rather than a complete compensation (Cressman & Henriques, 2009, 2010). In addition, the amount of recalibration was similar for both gradual and abrupt visuomotor perturbations (Cressman & Henriques, 2009, 2010; Salomonczyk et al., 2012).

Post-experiment questioning suggested that for group 3, the conflict between visual and proprioceptive targets was consciously noticed by 44% of the participants, compared to only 17% of group 1. We previously hypothesized that a fast gradual perturbation is less likely to be consciously perceived as it is more life-like and the abrupt group would behave similarly (Babu et al., 2023). Work from motor adaptation literature is inconsistent on how abrupt vs gradual perturbations affect learning but it has been suggested that an abrupt perturbation is more explicit, and the amount of learning differs from the implicit learning that results from a gradual perturbation (Buch et al., 2003; Kagerer et al., 1997; Saijo & Gomi, 2010; Schlerf et al., 2012; Werner et al., 2014). Though the cue conflict in the current study was often consciously perceived, this did not lead to reduced recalibration compared to group 1. This is consistent with previous studies (Hsiao et al., 2022; Hsiao & Block, 2024; Modchalingam et al., 2019), and with the idea that noticing the conflict does not override intrinsic belief in a common cause.

It is interesting that all of the change in performance for group 3 happened immediately upon introduction of the cue conflict, with no further recalibration while the perturbation was held constant at 70 mm. This is consistent with our previous finding that after a gradual mismatch increased to 70 mm at various rates, additional trials at a constant 70 mm cue conflict did not cause more recalibration (Babu et al., 2023). The proprioceptive recalibration observed in a motor adaptation study with an abrupt 30^°^ perturbation was similarly quick, reaching full magnitude after 6 trials with no effect of additional trials (Ruttle et al., 2016). Ruttle and colleagues demonstrated a similar time course of recalibration with an exposure paradigm where subjects moved their hand actively or passively along a channel that deviated from the cursor (Ruttle et al., 2018). The literature on proprioceptive recalibration thus seems consistently to indicate that it happens quickly, and then remains constant if the cue conflict is held constant. This is markedly different from motor adaptation, which would be expected to continue as long as systematic errors associated with the movement are detectable. For this reason the term “adaptation curve” is often used to describe motor adaptation, as error reduction continues for many trials even in the presence of a constant perturbation (Smith et al., 2006).

### Effect of feedback on visuo-proprioceptive recalibration: Groups 2 and 4 vs. 1 and 3

Feedback in a cue-conflict paradigm adds the possibility of systematic movement errors that lead to motor adaptation. Motor adaptation involves updating movements to account for internal and external changes. The brain generates a prediction of movement outcome to compare with the actual sensory consequences; a consistent discrepancy leads to updates in the model that generates predictions (DeSouza et al., 2000; Miall et al., 2007).

In the present study, the initial response to the perturbation shows significant differences between the two abrupt groups, 3 and 4, for visual and proprioceptive performance. This suggests that when there is an abrupt perturbation, the effect of feedback is immediately evident on exposure to the cue conflict. In contrast, with a gradual perturbation the effect of feedback (group 1 vs. 2) was more evident during the perturbation. The presence of feedback in general increased proprioceptive target overshoot and decreased visual target undershoot, which we predicted if the indicator hand is undergoing visuomotor adaptation. In other words, since the visual target is consistently further away than expected due to forward offset, the brain should interpret this as a sensory prediction error requiring the motor command to move the indicator hand a larger distance forward. Unlike visual and proprioceptive recalibration, which should shift indicator finger endpoints on the two unimodal target types closer together, motor adaptation would shift all indicator finger endpoints further forward.

Because pointing task performance in the feedback groups reflects motor adaptation in the indicator hand, we included the ruler task as an alternative method to assess proprioceptive recalibration in a way that did not involve the indicator hand. We predicted that if proprioceptive estimates by the ruler task showed greater recalibration in the feedback groups than the no-feedback groups, it would mean that the presence of sensory prediction errors increased proprioceptive recalibration beyond what would happen only in response to the cue conflict (Rossi et al., 2021). This would be analogous to what is thought to happen in a visuomotor cursor rotation task: proprioceptive recalibration is driven both by the cue conflict and the sensory prediction errors. Of course, in the cursor rotation paradigm, the cue conflict and the sensory prediction errors both pertain to the hand performing target-directed reaches. In contrast, in the present task, the target hand is subject to a cue conflict while stationary and sensory prediction errors are directly related to the indicator hand. The situation is ambiguous nonetheless; the indicator finger movements are aimed at the perceived target position, so the sensory prediction errors are not fully divorced from the target hand.

The ruler task results were neither fully inconsistent with nor fully consistent with our prediction. For the feedback groups only (2 and 4), the proprioceptive target pointing task yielded significantly more change in behavior than the ruler task, and there was no between-group difference on the ruler task. This is consistent with feedback (and sensory prediction errors) not affecting proprioceptive recalibration. On the other hand, there were significant pre-post changes in the ruler task only for the feedback groups, 2 and 4, which would be consistent with greater proprioceptive recalibration associated with feedback.

It should be noted that the ruler task may not be a sufficiently sensitive measure to detect a true between-group difference in response to feedback. In addition, the ruler task can be considered a high-level proprioceptive task, as the judgement involves a different frame of reference compared to the low-level pointing task which involves only making comparisons with one’s own body (Héroux et al., 2022; Wali & Block, 2024). To conclusively answer this question, future work will need to combine more than one method of inducing recalibration with more than one method of measuring it. These results underline the importance of broadening the range of tasks and assessments to better understand how narrowly (or broadly) a change in perception affects other perceptual and motor processes.

### Visual recalibration and effort minimization: Groups 5 and 6

In the absence of feedback, we interpret undershoot of visual targets as a reflection of visual recalibration (perceiving the visual cue as closer to the proprioceptive cue than it really is). While previous experiments have shown both visual and proprioceptive recalibration (Block & Bastian, 2011; Rand & Heuer, 2019), there are some paradigms that show little to no visual recalibration (Simani et al., 2007; Zbib et al., 2016). These paradigms do not necessarily measure visual recalibration explicitly, but rather infer the absence of visual recalibration based on inter-limb transfer (Clayton et al., 2014; Mostafa et al., 2014). The presence of visual recalibration also likely depends on the relative weighting of the visual cue, which is highly task-dependent. In our previous use of the current paradigm (no-feedback version), people tend to rely more heavily on proprioception than vision (Block & Sexton, 2020), and proprioception is also thought to be more heavily weighted for some computations than others (Lateiner & Sainburg, 2003; Mon-Williams et al., 1997; Sarlegna & Sainburg, 2006; Sober & Sabes, 2003; van Beers et al., 2002). Recalibration is thought to be inversely related to weighting, so relying more on proprioception than vision would be expected to result in a larger amount of visual recalibration (Ghahramani et al., 1997). Task-relevant, high-salience visual feedback as given in many cursor rotation studies is likely to cause people to rely much more on vision, which could substantially reduce visual recalibration. Giving the cursor feedback on a different horizontal plane, however, likely reduces reliance on visual cues and could increase visual recalibration (Rand & Heuer, 2019).

In groups 1-4, the cue conflict was always created by shifting the visual cue forward (away from the subject) relative to the proprioceptive cue. This raises the question of whether visual cue undershooting could instead be a result of participants wishing to minimize the indicator hand’s movement effort. With no feedback about the indicator hand, the participant receives no negative consequences for doing so, or incentives for not doing so. The positioning of the two tactile markers and the 5 potential indicator finger start positions is meant to keep everything well within arms’ reach, even with a 70 mm forward offset. In any case, to test this possibility, subjects in two additional groups performed reaches to a visual cue that moved away gradually, with (group 5) or without (group 6) intermixed proprioceptive trials. We hypothesized that with no bimodal trials to create a cue conflict (group 5) or no non-visual trials at all (group 6), there would be no visual recalibration and therefore subjects would closely follow the visual cue without undershooting it. As predicted, subjects were able to point at the visual cue throughout the 0 – 70 mm shift in groups 5 and 6. Group 1 had significantly more visual undershoot (recalibration) than group 5 or 6, which suggests that the undershoot varies as a consequence of the cue conflict arising in the paradigm, and not due to effort minimization.

Visual target undershoot was not significantly different from zero for Group 6, meaning that subjects’ pointing accuracy on the visual target did not change even when the target moved 70 mm away. It is interesting that in Group 5 there was still some undershoot of the visual target even though there were no bimodal targets during block 2. This might indicate that the veridical block 1, which did have bimodal targets, was sufficient for subjects to associate the visual and proprioceptive cues as both being associated with their hand, even though not presented simultaneously. It is has been found that when two cues are spatially close to each other and their appearance is statistically correlated they are perceived as arising from the same underlying cause, allowing them to interact and influence perception (Hairston et al., 2003; Körding et al., 2007; Wallace et al., 2004).

## Conclusions

The present results suggest that visuo-proprioceptive recalibration occurs with similar magnitude whether the cue conflict is introduced abruptly or gradually. However, the timing differs, with recalibration happening all at once when an abrupt conflict is imposed. The presence of performance feedback influenced motor adaptation in the indicator hand, but recalibration in the target hand proceeded regardless. Error feedback may play a smaller role in recalibration for a static hand as in the present study, than for a moving hand as in the visuomotor adaptation literature.

## Acknowledgement

Funding was provided by National Institute of Neurological Disorders and Stroke (Grant No. R01 NS112367 to HJB).

